# Does flavor-nutrient learning promote or protect against diet-induced obesity? Individual differences in conditionability predict resistance to weight gain in rats

**DOI:** 10.64898/2026.04.12.718046

**Authors:** Kevin P. Myers

## Abstract

Flavor-nutrient learning (FNL) refers to learning associations between a food’s flavor and the rewarding appetition signals that arise from post-oral nutrient sensing during or after a meal. In rodent models FNL reliably produces strong flavor preferences and increased intake of nutrient-paired flavors, implicating FNL as a presumptive obesogenic influence in the modern environment. However, evidence that FNL plays a causal role in diet-induced obesity is ambiguous. We have previously shown that degree of weight gain on a high-fat/sugar diet is associated with stronger FNL responses, but direction of causation was unclear. This paper reports three experiments investigating whether individual differences in FNL ‘conditionability’ are linked to obesity proneness prior to obesity onset. Two experiments comparing selectively-bred obesity-prone vs resistant strains found no strain differences in FNL. A third study in lean, outbred rats evaluated whether baseline individual differences in FNL prospectively predict weight gain on a cafeteria diet. Unexpectedly, rats who showed the strongest learned increase in intake of a nutrient-paired flavor subsequently gained the least weight when switched to cafeteria diet, suggesting FNL protects against weight gain. In fact, individual differences in FNL explained a portion of variance in cafeteria weight gain over and above measured kcal intake, implying a function for FNL in adaptively modulating metabolic responses to energy intake. Collectively, several studies have now shown individual differences in obesity proneness to be either positively correlated, uncorrelated, or negatively correlated with FNL, calling for a more nuanced view of how appetition influences intake and energy balance.

## 1. Introduction

Food choice, meal size, and meal patterning are strongly influenced by learned associations between a food’s sensory properties and its postingestive nutritional and metabolic impacts. Through this process of flavor-nutrient learning (hereafter, “FNL”) a food’s flavor becomes a Pavlovian conditioned stimulus (CS) associated with the unconditioned stimuli (US) generated by post-oral nutrient sensing, leading to conditioned changes in flavor evaluation.

Extensive work studying FNL in rodent models has used the ‘electronic esophagus’ approach pioneered by Sclafani (e.g., Elizalde & Sclafani, 1990; Lucas & Sclafani, 1989; Sclafani & Nissenbaum, 1988), which enables experimental control over the flavor-nutrient relationship unconfounded by the taste of the nutrient itself. Animals are trained with a flavor (CS+) accompanied by intragastric (IG) nutrient infusion, and a different (CS−) flavor accompanied by IG water. In CS+ flavor sessions, as in an ordinary meal, the peripheral signals arising from post-oral nutrient sensing that stimulate central reward circuitry are referred to as *appetition signals*. In contrast to satiation signals (gastric stretch, CCK, GLP-1, etc.) which bring the meal to an end, appetition signals are positive, feed-forward signals that increase intake and enhance CS+ flavor evaluation (Sclafani, 2013; Sclafani et al., 2026; Zukerman et al., 2013).

The immediate (unconditioned) and subsequent (conditioned) changes in behavioral responses to the CS+ flavor include at least three *appetition effects* (see reviews by Berthoud et al., 2021; Myers, 2018). 1) The *immediate appetition effect* is seen when post-oral appetition signals generated within a CS+ session stimulate intake within that session. 2) A *conditioned acceptance effect* is seen in subsequent CS+ sessions when intake rate is higher from the start. 3) Finally, a *conditioned preference effect* is reliably observed, whereby after training, animals strongly prefer the CS+ flavor over CS− in a two-bottle choice. Importantly, after training, conditioned CS+ acceptance and preference effects are evident even when the flavor is no longer accompanied by nutrient infusion, as they reflect learned associations between the flavor and rewarding appetition signals. In this paper, the term flavor-nutrient learning (FNL) refers collectively to these three behavioral changes, although they may individually play different roles in food intake and energy balance.

A notable attribute of FNL is that all three appetition effects promote increased caloric intake in the short term, by steering food selection (conditioned preference) and increasing meal size (immediate appetition and conditioned acceptance). Those responses would assist foraging animals to maximize energy intake, but could promote overconsumption in the modern obesogenic environment. However, whether FNL actually acts to promote long-term positive energy balance and weight gain has never been clearly established. Some early ‘electronic esophagus’ studies provide mixed support (Sclafani & Nissenbaum, 1988; Sclafani et al., 1996). Other studies suggest FNL may protect against weight gain, as rats gain more weight when flavor-nutrient relationships are inconsistent (Warwick & Schiffman, 1991). Thus, while ample evidence shows FNL increases short-term energy intake (Berthoud et al., 2021; Myers, 2018; Sclafani, 2013), whether FNL actually plays a role in diet-induced obesity is far from clear.

We have previously reported initial evidence that diet-induced obesity is positively associated with sensitivity to FNL (Wald & Myers, 2015). When fed an obesogenic high-fat, high-sugar (HFHS) diet, rats who gained the most weight subsequently showed stronger FNL compared to rats who gained less weight. But this association does not establish direction of causation. Differences in FNL ‘conditionability’ could be a pre-existing factor in obesity proneness, or could result from long-term impacts of having consumed more HFHS diet.

The issue is further complicated by noting that the only other published rodent study linking FNL and diet-induced obesity (Woods et al. 2016) observed *impaired* FNL in obese rats, contrary to the findings of Wald & Myers (2015). These different outcomes may stem from important procedural differences, as Woods et al (2016) used an alternative FNL protocol initially developed by Warwick & Weingarten (1994), which does not involve IG nutrient infusion. Instead, the CS+ and CS− flavors are consumed in vehicle solutions of similar palatability but different energy density (i.e., CS+ in glucose versus CS− in dilute glucose + saccharin). This “oral training” protocol, which we have also used successfully in our lab for some studies (Palframan & Myers, 2016), is valued for its technical simplicity, but there may be important yet unrecognized differences in FNL responses produced by these two methods.

The apparently contradictory outcomes between our previous findings (Wald & Myers, 2015) and Woods et al (2016) indicate, at a minimum, that the links between FNL and obesity are complex and require further investigation. Conclusively testing the hypothesis that FNL causally promotes obesity would require gain- or loss-of-function manipulations with high specificity for FNL over a relatively long term to test whether experimentally enhancing or attenuating FNL increases or prevents weight gain. Since the neural circuity underlying FNL is not fully characterized, in the absence of validated methods with high functional specificity, the present experiments seek corroborating evidence that is consistent or inconsistent with the causal hypothesis. These experiments are based on the rationale that if FNL does play a causal role in diet-induced obesity, it would be expected that *individual differences* in FNL measured prior to any obesogenic diet exposure would be linked to differential obesity proneness. Therefore, the current experiments first used selectively-bred obesity prone (OP) and obesity-resistant (OR) rat strains, and assessed FNL prior to any exposure to an obesogenic diet. Experiment 1 used IG training (after Wald & Myers, 2015) and Experiment 2 used oral training (after Woods et al., 2016). If selectively-bred rats known to be obesity prone (but not actually obese) demonstrate stronger FNL, that would be corroborative evidence for FNL as an obesogenic risk factor.

Experiment 3 further investigated whether sensitivity to FNL confers obesity-proneness or resistance by testing a large group of outbred rats to determine whether individual differences in FNL would prospectively predict subsequent weight gain. A large sample of outbred Sprague-Dawley rats was initially assessed for sensitivity to FNL before being given extended access to an obesogenic cafeteria diet to determine if rats who initially showed the stronger or weaker FNL responses gained different amounts of weight.

## 2. Experiment 1

Experiment 1 was designed to test whether selectively-bred obesity-prone versus obesity-resistant rats show differential sensitivity to FNL when tested in young adulthood prior to any obesogenic diet exposure. Rats from the Levin DIO and DR strains (originally derived from a Sprague-Dawley background, Levin et al., 1997) were used, although here the designations “OP” and “OR” are used (Vollbrecht et al., 2015) to convey that when studied prior to any obesogenic diet exposure the two strains are both lean and remain similar in weight on a standard grain-based rodent chow. The FNL training protocol closely mimicked our previous study assessing FNL in diet-induced obesity (Wald & Myers, 2015) by using an IG nutrient infusion (5 ml of 6% glucose) that produces only a moderate degree of FNL, to avoid floor and ceiling effects in the OP vs OR strain comparison.

### 2.1 Experiment 1 Methods

For all experiments, animal procedures were approved by the Bucknell University IACUC and were performed according to the NIH Guide for the Care and Use of Laboratory Animals, 8^th^ edition. Rats were housed individually in polycarbonate cages with corn cob bedding and wire lids, in colony rooms maintained at 20-26°C with a reversed 12:12 light:dark cycle. All behavioral procedures were performed during the dark phase. Unless otherwise specified rats were fed standard grain-based chow (TestDiet 5001). When food restricted, rats were fed daily rations sufficient to maintain 90-95% of their free-feeding weight, given at least 1 hr after behavioral testing daily.

Subjects in Experiment 1 were 30 adult female rats (15 OP and 15 OR) born in our colony to DIO- and DR-strain breeding stock graciously donated by Dr. Carrie Ferarrio. Females were used to be consistent with our prior study (Wald & Myers, 2015). Rats were ∼65 days old at the outset and the two strains did not differ in body weights (Mean ± SD for OP and OR were 207.0 ± 11.1 g and 202.2 ± 11.6 g). Each rat had a Silastic IG catheter surgically installed into the fundus of the stomach. The catheter ran subcutaneously from the abdomen to the nape of the neck where it terminated in a Luer-Lock connecter which remained capped when not in use. Surgeries were performed with isoflurane anesthesia. Rats were treated postoperatively with carprofen (10 mg/kg SC) and enrofloxacin (10 mg/kg daily SC) immediately after surgery and the next two days. By the third postoperative day all rats met or exceeded their pre-operative body weight. They remained on ad libitum chow for at least 8 days, at which time they were fed restricted rations (10-11 g/day) daily after training/testing sessions as described below.

#### 2.1.1 Apparatus

Flavor-nutrient conditioning sessions were conducted in 10 identical acrylic chambers (25 cm L X 25 cm W X 32 cm H) with a stainless steel wire grid floor. For each session a rat’s IG catheter was attached to infusion tubing suspended overhead on a standard fluid swivel/counterbalance arm, connected to a computer-controlled syringe pump. The drinking bottle was held on a motorized bottle retractor (modified Med Associates ENV-252) and drinking was monitored by electronic contact lickometers. In every session, onset of each rat’s IG infusion was triggered by the first sustained licking bout and consisted of 5 ml infused at a constant rate of 0.33 ml/min for 15 min. This ensured that all rats received identical IG infusions irrespective of individual differences in baseline intakes or licking rate. The bottle remained available for the remainder of the 30-min session and total intake was measured by weight.

#### 2.1.2 Conditioning

Prior to conditioning rats were first acclimated to daily 30-min drinking sessions in the apparatus, with water infused 5 ml/15 min while consuming plain 0.1% saccharin. The baseline CS− sessions occurred for the next three sessions, in which the 0.1% saccharin was now flavored with either cherry or grape Kool-Aid powder (0.05%, counterbalanced across rats) to establish baseline intake of CS− with IG water infusion. In the next three sessions rats received their CS+ flavor (the opposite Kool-Aid flavor) with the same session parameters except that 6% glucose (5 ml/15 min) was infused instead of water. Finally, because immediate flavor recency could bias subsequent preference and we have anecdotally observed non-specific elevation of solution intakes in the 1-2 days immediately after glucose-infused sessions, three additional sessions consisting only of 0.1% saccharin + novel strawberry Kool-Aid flavor, with no IG infusions, were conducted before further testing.

#### 2.1.3 Post-conditioning testing

Rats were tested for conditioned acceptance effects on two consecutive days in the lickometer boxes with 30-min access to a single bottle containing either CS+ or CS− flavor in 0.1% saccharin, in counterbalanced order, with no IG infusion. Intakes were measured to the nearest 0.1 g by weight.

Two-bottle tests for conditioned preference were conducted in the home cages instead of the lickometer apparatus, because spout position was not alternated during the training sessions (only the right side of the bottle retractor was used to minimize variability in licking rates during the critical training sessions), meaning strong side preferences would be likely to bias two-bottle tests in the lickometer apparatus. Rats were familiarized with home cage two-bottle choice tests for two sessions with 0.1% saccharin + 1% fructose versus 0.05% saccharin + 0.5% fructose, with left/right bottle placement reversed daily. Then rats were given 30-min/day two-bottle choice tests with CS+ vs. CS− flavors (i.e., grape vs. cherry) in 0.1% saccharin, repeated on two consecutive days with the left/right position of the flavors alternated. Intakes (measured by weight) of each flavor were averaged across the two repetitions.

### 2.2 Experiment 1 Results

Three rats (two OR and one OP) were removed mid-experiment because of problems with their IG catheters, for a final n = 13 OR and 14 OP. Differential sensitivity to FNL was assessed in several ways. The immediate appetition response would be indicated by elevated licking rate within the first CS+ (IG glucose) session relative to the preceding CS− (IG water) session. Conditioned acceptance (increased intake) would initially be evident in the progressive elevation of licking across the three consecutive CS+ sessions. Next, greater intake of CS+ than CS− in subsequent one-bottle intake tests without IG infusions demonstrates this is a conditioned response to the CS+ flavor. Finally, conditioned preference for CS+ over CS− was determined by the home-cage two-bottle choice tests.

#### 2.2.1 Immediate appetition effects

Cumulative licking during the 30-min conditioning sessions is depicted in Figure 1A and 1B. As expected, all rats demonstrated immediate appetition by consuming more in the first CS+session than the preceding CS− session, reflecting the rapid intake-stimulating effect of IG glucose (repeated measures ANOVA on total licks in those two sessions only, main effect of CS− vs CS+, F (1,25) = 23.82, *p* < .001.) Regardless of session type, OP rats always consumed more than OR rats overall (main effect of Strain, F (1,25) = 6.23, *p* = .020). However, there was no strain difference in immediate appetition, as the increase from CS− to CS+ was proportionally similar for both strains (CS X Strain interaction, F(1,25) = 0.07, *p* = .78)

**Figure 1.**
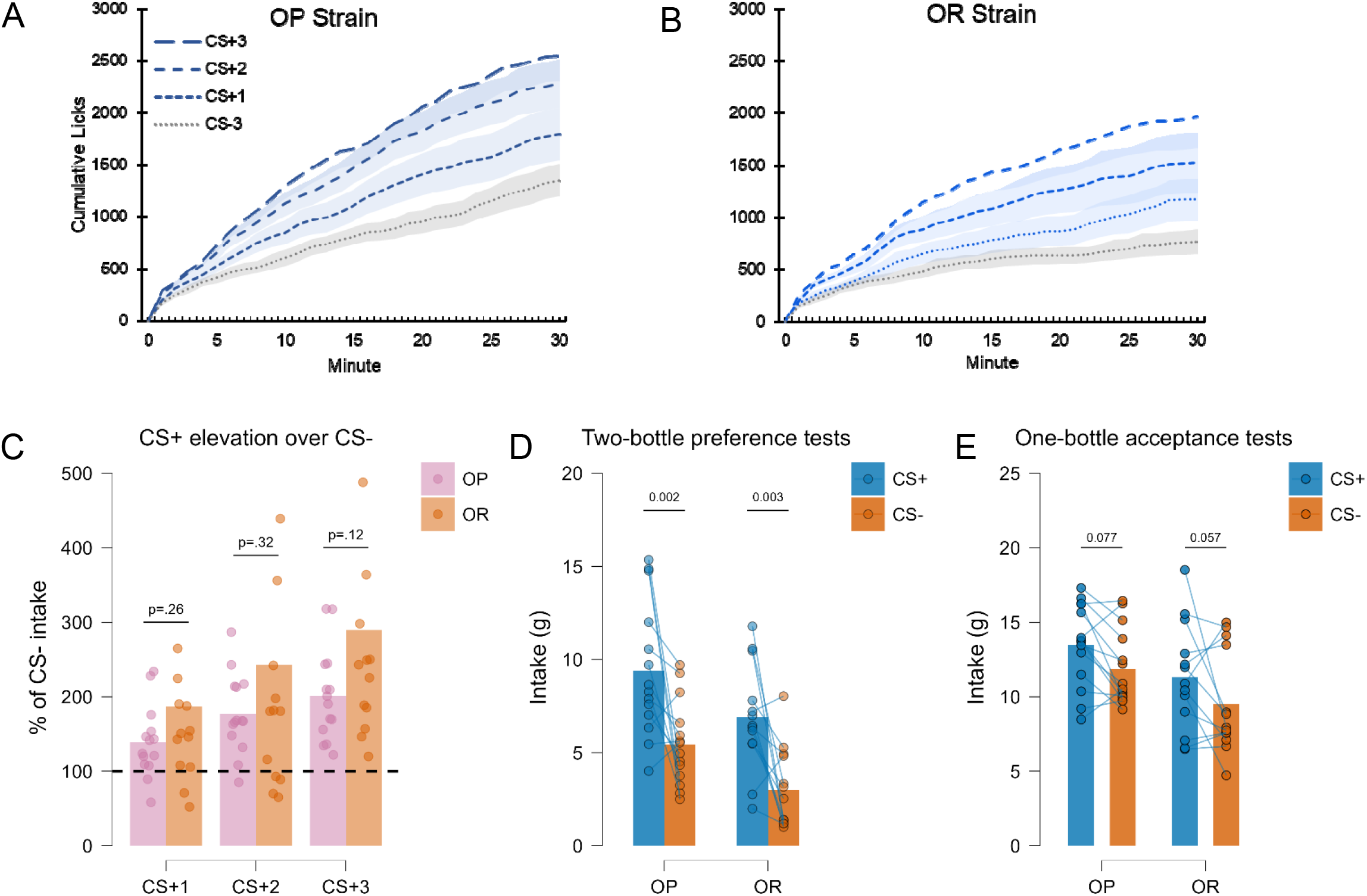
(A&B) Cumulative licks by OP rats (A) and OR rats (B) in the last CS-training session (flavor paired with IG water) and each of the three CS+ sessions (flavor paired with IG glucose). (C) 30-min intakes during each of the three consecutive CS+ conditioning sessions expressed as percentage relative to the final CS− session for OP and OR strain rats. (D) Intakes of OP and OR rats in the post-conditioning 30-min two-bottle preference tests between CS+ and CS− flavors in the absence of IG infusions. (E) Intakes of OP and OR rats in 30-min one-bottle acceptance tests with the CS+ and CS− flavors in the absence of IG infusions. All *p* values are two-tailed, Bonferroni-corrected pairwise tests.

#### 2.2.2 Increased acceptance across conditioning sessions

Progressively increasing acceptance across CS+ training sessions (Figures 1A and 1B, and expressed as CS+ intake as percentage increase over CS− baseline, Figure 1C) was also similar in both strains. By the third CS+ session intakes increased to 289% ± 54.2 (Mean ± SEM) over CS− in OR rats and 202% ± 16.6 in OP rats. While nominally larger in OR rats, the strain difference was not statistically significant (Figure 1C, repeated measures ANOVA, main effect of Session, F (1,25) = 19.29, *p* < .001, no main effect of Strain, F (1,25) = 1.63, *p* = .21, no Session X Strain interaction, F (1,25) = 1.16, *p* < .32).

#### 2.2.3 Post-conditioning preference and acceptance tests

Two-bottle tests in the home cage revealed all rats had acquired a strong learned CS+ preference and this did not differ between OP and OR strains. Rats consumed significantly more CS+ than CS− in the two bottle tests (Figure 1D, repeated measures ANOVA main effect of Flavor, F(1,25) = 22.55, *p* < .001) and although total consumption was greater in OP rats (main effect of Strain, F(1,25) = 12.32, *p* = .002) the relative CS+ preference did not differ by Strain (Flavor X Strain interaction, *F*(1,25) < .001, p = .99).

Post-conditioning intakes in separate one-bottle sessions without IG infusions demonstrated a relatively weak but statistically significant conditioned acceptance effect, which once again did not differ between OP and OR strains. Rats consumed significantly more in CS+ sessions than CS− sessions (Figure 1E, repeated measures ANOVA, main effect of Flavor, F (1,25) = 7.38, *p* = .018). Once again, OR rats consumed more of both flavors than OP rats (main effect of Strain, F (1,25) = 4.60, *p* = .04) but the increased CS+ intake relative to CS− was proportionally similar between strains (Flavor X Strain interaction, F (1,25) = 0.03, *p* = .88).

### 2.3 Experiment 1 Summary

Contrary to expectations, the obesity prone and resistant strains did not differ in any measure of flavor-nutrient learning. The only consistent strain difference was that OP rats’ intakes of all solutions were generally higher. But taking those baseline differences into account, the immediate appetition response to IG glucose, the learned increases in CS+ meal size, and the learned CS+ preference were all equivalent in the two rat strains. Therefore, in our previous study demonstrating a positive link between degree of obesity and FNL, the heightened FNL responses were more likely the result of obesity and/or the history of excess sugar and fat consumption by the obese rats, and not a pre-existing risk factor for obesity.

## 3. Experiment 2

Experiment 2 also investigated whether obesity-prone and resistant rats exhibit pre-existing differences in FNL, using an oral training protocol like that used in Woods et al. (2016). Since the only two previously published rodent studies of FNL and diet-induced obesity used different FNL protocols, finding consistent or inconsistent results across training methods could be informative as to whether these two experimental protocols truly rely on the same neurobehavioral mechanisms.

In Experiment 2, female OP and OR rats were trained in sessions alternating between a CS+ flavor in a 6.8% glucose solution and a CS− flavor in a solution of approximately equal palatability but lower energy density, specifically 1% glucose + 0.125% saccharin (hereafter, “G+S”). Those concentrations were selected by testing naive rats in a series of two-bottle preference tests between G+S versus glucose concentrations ranging from 5-10%. By interpolation, 6.8% glucose was predicted to be equally preferred to G+S, and that was confirmed with an additional group of naive pilot rats. Using these concentrations, during training sessions the CS+ flavor delivered 0.27 kcal/ml and the CS− only 0.04 kcal/ml.

### 3.1 Experiment 2 Methods

Subjects were 32 young adult female rats (16 OP and 16 OR), with similar average weights (OP = 214.0 ± 12.5 g, OR = 211.1 ± 10.0 g). Rats were maintained on restricted chow rations (∼11 g/rat/day) and were first familiarized with the daily 30-min sessions with a bottle containing 50ml of G+S flavored with a different novel Kool-Aid flavor (0.05% orange, blue-raspberry, lemon lime) each day to habituate flavor neophobia. The G+S bottle was given in the home cage for 2 hrs, followed 2 hrs later by chow rations.

Then flavor-nutrient conditioning sessions were conducted over 4 consecutive days, consisting each day of either CS+ flavor (.05% grape or cherry Kool-Aid powder) in 6.8% glucose or the opposite CS− flavor in G+S. Because OP rats typically consumed more G+S than OP rats in the initial G+S familiarization sessions, the CS+ and CS− were limited to 15 ml per session so that any eventual differences in FNL responses would not be attributable to differential consumption during training. The order of conditioning sessions was + − − + or − ++ − counterbalanced across rats. Each day, a bottle containing 15 ml of the CS+ or CS− solution was placed on the home cage approximately 2 hours after dark onset. On the first conditioning day, bottles were checked 2 hrs later, and any unfinished solution remained on the cage for another 1 hr, at which time chow rations were fed. On all subsequent days, all rats finished the entire 15 ml within 1 hr.

Following the 4-day conditioning procedure, rats were familiarized with two-bottle testing in the home cages, as in Experiment 1, using 1% glucose + 0.125% saccharin versus 0.5% glucose + 0.062% saccharin. Then conditioned preference for the CS+ over CS− flavor was measured in 30-min two-bottle choice tests with the CS+ and CS− flavors both given in the equivalent G+S vehicle. This test was repeated on two consecutive days with the left/right position reversed. Results are the average of the two tests.

### 3.2 Experiment 2 Results

#### 3.2.1 Conditioned Preference

All rats strongly preferred CS+ over CS− in the two-bottle choice, Figure 2A, Flavor X Strain repeated measures ANOVA, main effect of Flavor, *F*(1,30) = 220.78, *p* < .001. OP rats once again consumed more overall (main effect of Strain, *F*(1,30) = 13.57, *p* < .001) but the strains’ relative preference for CS+ over CS− did not differ, Flavor X Strain interaction, *F*(1,30) = 3.72, p = .063. Although the Flavor X Strain interaction was close to statistically significant, expressing preference strength as CS+ intake as a percentage of total intake and comparing between strains again confirmed no difference, Mean ± SEM for OP = 85.1% ± 2.91, OR = 87.4% ± 2.05, *t*(30) = 0.673, *p =* .51.

**Figure 2.**
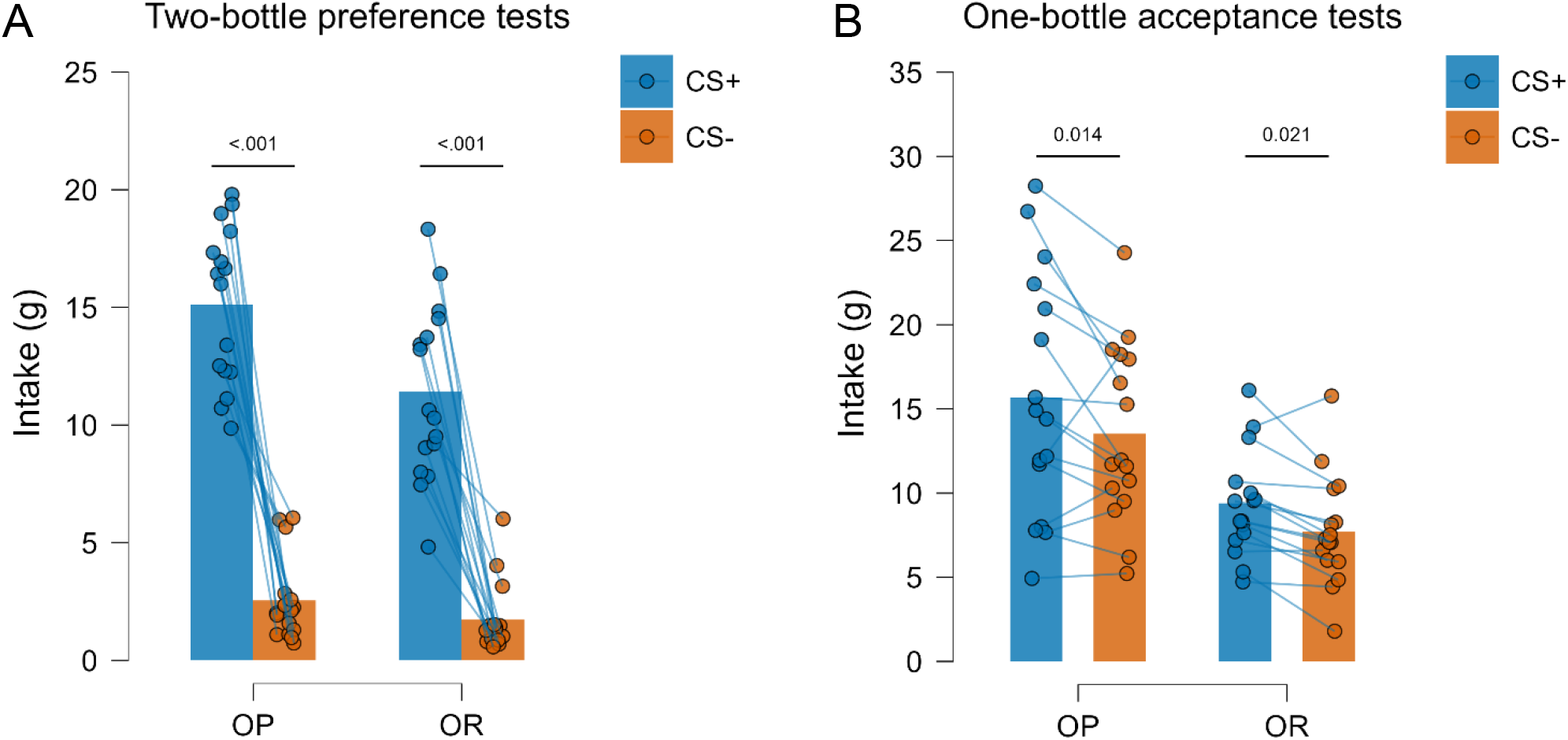
(A) Intakes of OP and OR rats in the post-conditioning 30-min two-bottle preference tests between CS+ and CS− flavors, now both in the equivalent G+S vehicle. (B) Intakes of OP and OR rats in 30-min one-bottle acceptance tests with the CS+ and CS− flavors, both in G+S. All *p* values are two-tailed, Bonferroni-corrected pairwise tests.

#### 3.2.2 Conditioned Acceptance

Similar conditioned acceptance was observed in both strains. Rats consumed significantly more in one-bottle CS+ sessions than CS− sessions, Figure 2B, Flavor X Strain repeated measures ANOVA, main effect of Flavor, *F*(1,30) = 7.38, *p* = .001. OP rats consumed more of each flavor than OR rats (main effect of Strain, *F*(1,30) = 12.90, *p* = .001) but the increased CS+acceptance relative to CS− was proportionally similar between strains (Flavor X Strain interaction, *F*(1,30) = 0.20, *p* = .66).

### 3.3 Experiment 2 Summary

Experiments 1 and 2 both observed no differences between selectively bred obesity-prone and resistant rats in any behavioral measure of FNL. Both strains acquired strong conditioned preferences for a glucose-paired flavor as seen in the post-conditioning two-bottle choice tests. In Experiment 1, the IG protocol allowed for more comprehensive assessment of appetition effects, because it provides cumulative licking records during training sessions. Those measures revealed that OP and OR rats responded with similar intake stimulation by post-oral nutrient sensing, as seen in the immediate appetition response to glucose infusion, and in proportionally similar increased intakes over repeated CS+ sessions, and, like Experiment 2, similar conditioned CS+ acceptance in the absence of IG infusion.

Experiment 2 differed from Experiment 1 in using an alternative oral training protocol (Warwick & Weingarten, 1994; C. A. Woods et al., 2016) instead of IG infusions. There were also some procedural differences such as fewer CS+ sessions (two instead of three) and equating CS+ and CS− intake during training sessions. In spite of these procedural differences, Experiments 1 and 2 produced highly similar results.

Although these experiments do not strictly test the hypothesis that FNL acts to promote diet-induced obesity (which would require either gain- or loss-of-function manipulations to test whether experimentally enhancing or attenuating FNL alters weight gain) they are nonetheless inconsistent with that view, which would predict that sensitivity to FNL is part of the obesity-prone phenotype. The OP rats are known to be highly susceptible to obesity, as this strain has been selectively bred for rapid weight gain in response to an obesogenic diet. Yet they were not more ‘conditionable’ in FNL than their obesity-resistant counterparts. At the least, these experiments demonstrate that the behavioral and physiological differences that confer the obesity-prone phenotype in OP rats do not involve differences in the mechanisms of FNL.

## 4. Experiment 3

To further investigate whether FNL may play a causal role in diet-induced obesity, Experiment 3 was designed to determine if inherent individual differences in FNL can prospectively predict subsequent weight gain. A large group of young adult female rats (outbred Sprague-Dawley) was screened on several measures of FNL using the oral training method. Then all rats were switched to a palatable, energy-dense cafeteria diet comprising ad libitum chow plus a varied selection of palatable ultra-processed foods. If FNL is a causal factor in diet-induced obesity, one would expect that rats who showed the strongest FNL in the conditioning protocol would also gain the most weight when transitioned to a cafeteria diet. The cafeteria diet used was designed as a diet that would allow for FNL to operate. That is, the varied foods all had distinctive flavors and sensory properties and also different energy densities and macronutrient compositions, which should enable rats to learn about relationships between flavors and postingestive nutritive consequences.

Because the rationale for this study depends on precise measurement of individual differences in FNL conditionability at baseline, the parameters of FNL training were designed to target only moderately strong appetition effects to avoid both ceiling and floor effects that would obscure individual differences. In several unpublished pilot studies, we have observed that with the oral conditioning procedure, conditioned preference for CS+ over CS− is acquired very rapidly with minimal training, i.e., a single 30-min session with each flavor, consistent with similar published observations using the IG infusion protocol (Ackroff et al., 2009; Myers, 2007) whereas conditioned acceptance takes more training to emerge. This could mean that adequate training to detect individual differences in conditioned acceptance might produce ceiling effects in conditioned preference. To remedy this, rats underwent two different rounds of conditioning to assess conditioned preference and conditioned acceptance separately. In the first round, rats were trained with a single 30-min session with each CS, with grape or cherry as the CS+ flavor in glucose and the opposite CS− flavor in G+S, and conditioned preference for grape vs cherry was then measured in a two-bottle choice (both flavors now in G+S). Then, rats were re-trained with novel CS+ and CS− flavors (strawberry and peach-mango). The second round involved a single extended session with each CS, providing 100 ml of solution to consume overnight. After one session with each flavor, conditioned increase in acceptance for CS+ was measured in a series of one-bottle intake tests with unflavored G+S, CS+ in G+S, and CS− in G+S. This procedure allows a more precise measurement of both conditioned preference and conditioned acceptance to determine if either is a reliable predictor of subsequent weight gain.

### 4.1 Experiment 3 Methods

Subjects were 34 young adult female Sprague-Dawley rats weighing 226 ± 23.6 g (Mean ± SD). They were initially acclimated to daily chow restriction (11 g/day) and familiarized with 30 min/day sessions with G+S, with a different novel Kool-Aid flavor each day (0.05% blue raspberry, orange, lemon-lime) to habituate neophobic responses to novel flavors. In a separate group of 24 rats drawn from the same colony (littermates of the Experiment 3 subjects), 6.5% glucose was identified as equally preferred to 1% glucose + 0.125% saccharin (‘G+S’), using procedures described in Experiment 2.

The first round of single-trail flavor nutrient conditioning occurred on two consecutive days, with a single 30-min CS+ session (.05% grape or cherry in 6.5% glucose) and a single CS−session (the opposite flavor in G+S) in counterbalanced order. Each day chow rations were given 2 hours after the session. Then two-bottle preference tests (each flavor in G+S) were conducted as described in Experiment 2.

Beginning one week later, rats were re-trained with two novel flavors for assessment of conditioned acceptance. In this phase rats were fed chow rations early in the day (∼10 AM) and then an extended conditioning session began in the late afternoon (∼4PM). One CS+ session and one CS− session occurred in counterbalanced order on consecutive days. Rats were given a bottle containing 100 ml of the CS flavor (.05% strawberry or peach-mango Kool-Aid) in either 6.5% glucose or G+S. The bottle remained on the cage overnight, and all rats consumed the entirety of the solution in both sessions. Beginning two days later, they were familiarized with receiving 30-min access to unflavored G+S prior to daily chow rations. Then, on three consecutive days, intakes were measured in 30-min one-bottle sessions, one session each with unflavored G+S, CS+ in G+S, and CS− in G+S (order across days counterbalanced) to measure the conditioned increase in CS+ acceptance.

Rats were then returned to ad libitum chow for 5 days prior to beginning a 24-day period of ad libitum cafeteria diet. The cafeteria diet involved 12 different highly palatable, energy-dense foods varying in sensory characteristics and macronutrient content (e.g., vanilla pudding, hot dog, chocolate cereal, fried pork rinds). Extensive details are provided in the Supplementary Methods. Each day rats were provided ad libitum portions of three different cafeteria foods plus chow. The cafeteria foods were rotated daily so that all foods occurred once in each 4-day cycle. Intakes of each cafeteria food and chow were measured daily for the first 4 days and last 4 days of the 24-day diet exposure, and weight gain was measured every 4 days.

### 4.2 Experiment 3 Results

For descriptive purposes, results of the flavor-nutrient conditioning are depicted in Figure 3. Rats strongly preferred the CS+ over CS− following the first single-trial conditioning procedure, paired *t*(33) = 11.94, *p* < .001. In the one-bottle acceptance tests after the second round of conditioning, rats consumed significantly more CS+ flavor in G+S than either unflavored G+S or CS− flavor in G+S (both *p* < .001, Bonferroni adjusted).

**Figure 3.**
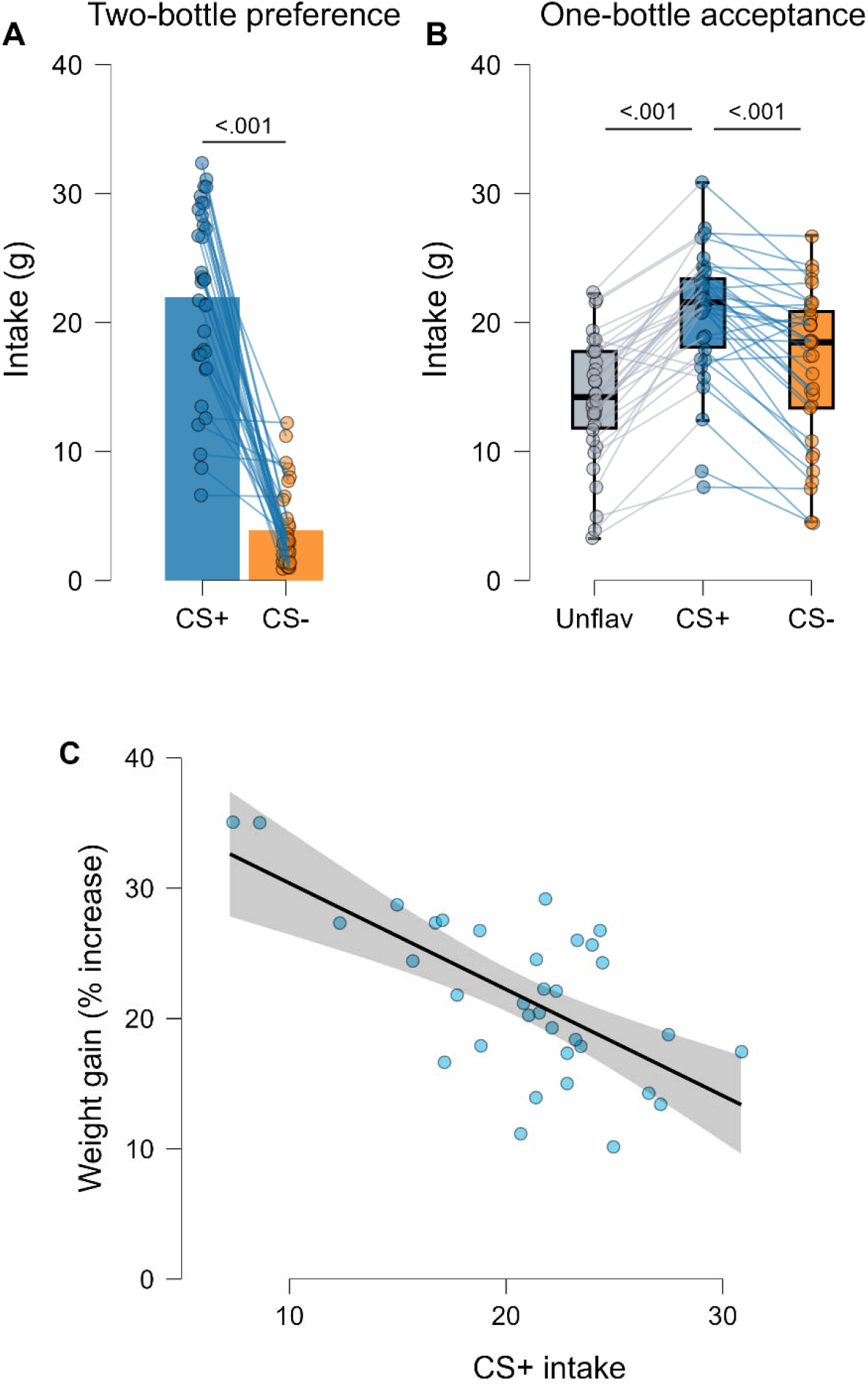
(A) Experiment 3 Intakes in the post-conditioning 30-min two-bottle preference tests between CS+ and CS− flavors, now both in the equivalent G+S vehicle. (B) Intakes in 30-min one-bottle acceptance tests with CS+ and CS− flavors, both in G+S. (C) Scatterplot depicting each individual rat’s cafeteria weight gain as a function of its CS+ intake in the one-bottle acceptance tests. Weight gain is the % increase in body weight after 24 days of cafeteria diet. All *p* values are two-tailed, Bonferroni-corrected pairwise tests.

To test the hypothesis that individual differences in FNL conditionability prospectively predicted subsequent cafeteria weight gain, multiple linear regression was used with percent increase in body weight across 24 days of cafeteria diet as the dependent variable and CS+ preference, CS+ acceptance, CS− acceptance, unflavored G+S acceptance, and baseline body weight as potential predictors.

Table 1 describes the preliminary pairwise correlation analysis. Conditioned CS+ preference strength in the two-bottle test was uncorrelated with weight gain or with conditioned acceptance measures. All intake measures in the one-bottle acceptance tests were correlated with weight gain, and were also correlated with each other (likely reflecting overall individual differences in attraction to sweetness and/or satiety sensitivity). The regression model began with all predictors entered and backward stepwise elimination with a removal criterion of *p* > 0.1. One-bottle CS+ acceptance emerged as the strongest predictor of weight gain and was the sole predictor retained in the final model (Table 2, *R^2^*= 0.42, *F*(1,33) = 24.6, *p* < .001). Rather unexpectedly, CS+ acceptance was *negatively* associated with weight gain, β = −0.66, *t*(33) = −4.96, *p* < .001, meaning that rats who showed the strongest tendency to increase intake of the glucose-paired CS+ flavor subsequently gained less weight on the cafeteria diet (Figure 3C).

**Table 1.** Correlations (n = 34) between weight gain and potential predictor variables in Experiment 3.

| Variable | *M* | *SD* | 1 | 2 | 3 | 4 | 5 | 6 |
| --- | --- | --- | --- | --- | --- | --- | --- | --- |
| 1. Cafeteria weight gain <sup>a</sup> | 21.7 | 6.2 | -- |  |  |  |  |  |
| 2. Two-bottle CS+ preference <sup>b</sup> | 83.6 | 13.8 | -.15 | -- |  |  |  |  |
| 3. One-bottle intake unflavored G+S <sup>c</sup> | 14.1 | 4.8 | -.45** | -.02 | -- |  |  |  |
| 4. One-bottle intake CS+ in G+S <sup>c</sup> | 20.7 | 5.0 | -.66*** | .11 | .73*** | -- |  |  |
| 5. One-bottle intake CS- in G+S <sup>c</sup> | 16.7 | 5.9 | -.46** | -.08 | .68*** | .78*** | -- |  |
| 6. Initial body weight | 322.5 | 26.7 | -.12 | -.06 | .04 | .10 | .03 | -- |

<sup>a</sup> Percentage increase, 24 days<sup>b</sup> CS+ intake as percentage of total intake in CS+ vs CS- two-bottle test<sup>c</sup> Intake (in g) of solution in 30-min one-bottle test\* $p < .05$ . \*\* $p < .01$ ., \*\*\* $p < .001$

**Table 2.** Multiple linear regression predicting cafeteria weight gain (n = 34)

| Model | Predictor | *B* | *SE* | 95%CI | $\beta$ | *t* | *p* |
| --- | --- | --- | --- | --- | --- | --- | --- |
| M<sub>0</sub> | Intercept | 47.11 | 12.22 | [22.08, 72.15] |  | 3.86 | < .001 |
|  | CS+ preference | -0.05 | 0.07 | [-0.19, 0.08] | -0.12 | -0.80 | .434 |
|  | One-bottle intake G+S | 0.12 | 0.27 | [-0.43, 0.69] | 0.10 | 0.45 | .654 |
|  | One-bottle intake CS+ in G+S | -0.65 | 0.32 | [-1.30, 0.01] | -0.53 | -2.03 | .052 |
|  | One-bottle intake CS- in G+S | -0.25 | 0.25 | [-0.76, 0.26] | -0.27 | -0.99 | .331 |
|  | Initial body weight | -0.02 | 0.03 | [-0.08, 0.05] | -0.07 | -0.51 | .616 |
| M<sub>4</sub> | Intercept | 38.54 | 3.50 | [31.41, 45.67] |  | 11.01 | < .001 |
|  | One-bottle intake CS+ in G+S | -0.82 | 0.16 | [-1.15, -0.48] | -0.66 | -4.962 | < .001 |

Notes: Backwards elimination using $p > 0.10$ for removalM<sub>0</sub> Adj. $R^2 = 0.37$ , $p = .003$ M<sub>4</sub> Adj. $R^2 = 0.42$ , $p < .001$

The ability of CS+ acceptance to predict weight gain was not an artifact of its correlation with sweet G+S intake, as the partial correlation between CS+ acceptance and weight gain while controlling for unflavored G+S intake was *r*(33) = −.57, *p* < .001, whereas the partial correlation between G+S intake and weight gain was only *r*(33) = −.06, *p* = .73. To further confirm that the predictive value of conditioned CS+ acceptance did not merely reflect some non-specific trait surrounding sweet solution intake or meal sizes generally, a second regression model was constructed to predict weight gain, entering unflavored G+S intake as the first predictor and then adding CS+ acceptance second. Unflavored G+S intake on its own was a significant predictor (Table 3, model M_1_, R^2^ = 0.20, F(1,33) = 7.99, *p* = .008). However, adding CS+ acceptance to the model explained a significant additional portion of the residual variance (Table 3, model M_2_, R^2^ change = 0.24, *F*_change_(1,31) = 13.05, *p* = .001).

**Table 3.**
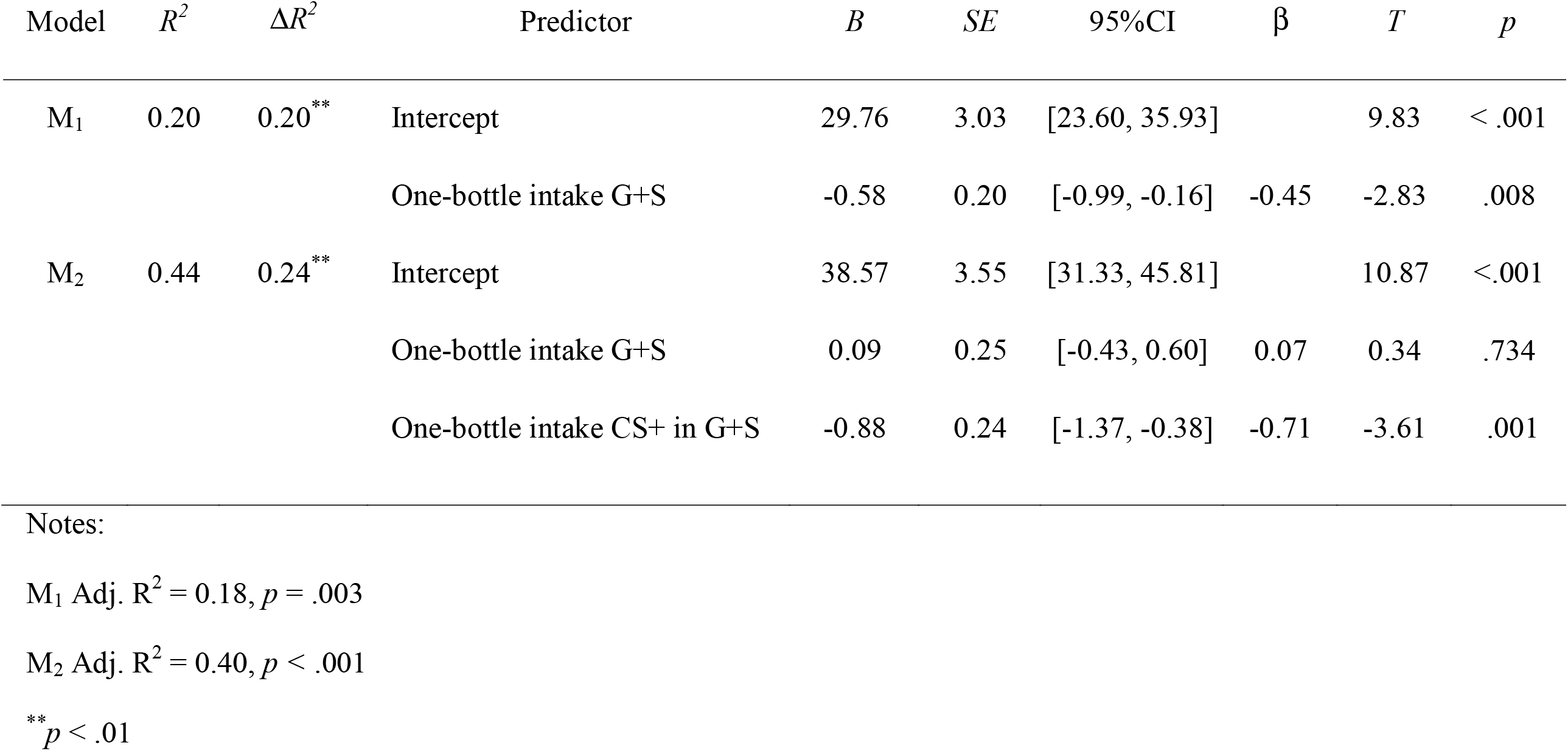
Multiple linear regression predicting cafeteria weight gain (n = 34) with unflavored G+S intake and CS+ acceptance.

| Model | $R^2$ | $\Delta R^2$ | Predictor | $B$ | $SE$ | 95%CI | $\beta$ | $T$ | $p$ |
| --- | --- | --- | --- | --- | --- | --- | --- | --- | --- |
| M<sub>1</sub> | 0.20 | 0.20<sup>**</sup> | Intercept | 29.76 | 3.03 | [23.60, 35.93] |  | 9.83 | < .001 |
|  |  |  | One-bottle intake G+S | -0.58 | 0.20 | [-0.99, -0.16] | -0.45 | -2.83 | .008 |
| M<sub>2</sub> | 0.44 | 0.24<sup>**</sup> | Intercept | 38.57 | 3.55 | [31.33, 45.81] |  | 10.87 | <.001 |
|  |  |  | One-bottle intake G+S | 0.09 | 0.25 | [-0.43, 0.60] | 0.07 | 0.34 | .734 |
|  |  |  | One-bottle intake CS+ in G+S | -0.88 | 0.24 | [-1.37, -0.38] | -0.71 | -3.61 | .001 |

Notes:
M<sub>1</sub> Adj. $R^2$ = 0.18, $p$ = .003
M<sub>2</sub> Adj. $R^2$ = 0.40, $p$ < .001
<sup>\*\*</sup> $p$ < .01

Finally, a third regression model further evaluated conditioned CS+ acceptance as a predictor of weight gain relative to a measure of actual caloric intake on the cafeteria diet. Total kcal intake measured for the first and last four days of cafeteria diet was entered first, and strongly predicted weight gain (Table 4, M_1_, *F*(1,33) = 11.03, *p =* .002). Adding the measure of CS+ acceptance explained a significant portion of residual variance (Table 4, M_2_, R^2^ change = 0.322, *F*_change_(1,31) = 23.27, *p* < .001). This indicates that individual differences in flavor-nutrient learning predicted weight gain independently of caloric intake, and that the combination predicted weight gain better than the kcal measurement alone.

**Table 4.** Multiple linear regression (n = 34) predicting cafeteria weight gain from kcal intake and CS+ acceptance.

| Model | $R^2$ | $\Delta R^2$ | Predictor | $B$ | $SE$ | 95%CI | $\beta$ | $T$ | $p$ |
| --- | --- | --- | --- | --- | --- | --- | --- | --- | --- |
| M<sub>1</sub> | 0.26 | 0.26<sup>**</sup> | Intercept | -6.60 | 23.15 | [-53.75, 40.56] |  | -0.29 | .778 |
|  |  |  | Kcal intake<sup>a</sup> | 0.09 | 0.03 | [0.04, 0.15] | 0.51 | 3.32 | .002 |
| M<sub>2</sub> | 0.58 | 0.32<sup>***</sup> | Intercept | 50.42 | 21.22 | [7.13, 93.70] |  | 2.38 | .024 |
|  |  |  | Kcal intake<sup>a</sup> | 0.09 | 0.02 | [0.04, 0.12] | 0.44 | 3.79 | <.001 |
|  |  |  | One-bottle intake CS+ in G+S | -2.30 | 0.47 | [-3.27, -1.34] | -0.57 | -4.87 | .001 |

Notes:
<sup>a</sup> Total kcal consumed in first four days and last four days of 24-day cafeteria diet
M<sub>1</sub> Adj. $R^2 = 0.23$ , $p = .002$
M<sub>2</sub> Adj. $R^2 = 0.55$ , $p < .001$
<sup>\*\*</sup> $p < .01$ , <sup>\*\*\*</sup> $p < .001$

### 4.3 Experiment 3 Summary

Unlike Experiments 1 and 2, which found no evidence linking FNL conditionability and obesity proneness when comparing OP vs OR rats, Experiment 3 found a novel and unexpected relationship in outbred rats. When tested in a FNL paradigm designed to detect individual differences in ‘conditionability’ by using relatively brief training with a moderate intensity nutrient US, a clear relationship was revealed between FNL and subsequent weight gain when rats were later transitioned to a palatable, energy-dense cafeteria diet. More specifically, conditioned acceptance, but not conditioned preference, negatively predicted weight gain. Rats who showed the largest stimulation of intake of the nutrient-paired CS+ flavor gained the least weight during 24-day cafeteria diet access. This association was robust, as conditioned acceptance was a stronger predictor of weight gain than baseline sweet taste intake or baseline body weight, and also increased the predictive accuracy of estimated kcal intake of the diet.

Not only is this the first evidence demonstrating that individual differences in FNL can prospectively predict weight gain, it is also notable that the association was opposite the direction that would be expected based on the simple view of FNL as a mechanism for promoting energy intake. However, it should be noted that the relationship found in Experiment 3 is inherently correlational. While not conclusively demonstrating that conditioned acceptance itself causes resistance to weight gain, it does indicate that conditioned acceptance is a behavioral marker of a resistance. The factors responsible for individual differences in conditioned acceptance are apparently unrelated to the phenotypic traits that differentiate selectively bred OP and OR rats, who showed no inherent differences in FNL.

## 5. General discussion

These experiments address the putative links between flavor-nutrient learning and diet-induced obesity, building on previous work that has shown both enhanced and impaired FNL responses after obesity onset (Wald & Myers, 2015; Woods et al., 2016). The current experiments were all based on the premise that individual differences in conditionability may be useful for identifying the functional relationship between FNL and weight. Other domains of Pavlovian conditioning (aversion, fear, evaluative conditioning) are marked by stable, moderately heritable individual differences in conditionability (Elkins, 1986; Hettema et al., 2003; Shumake et al., 2014) associated with real-world behavioral outcomes such as gambling (Brunborg et al., 2011) anxiety disorders (Orr et al., 2000; Wegerer et al., 2013) and cue reactivity in substance use (Glautier et al., 2000). If FNL plays any causal role in weight gain, then individual differences in the two should be associated.

Contrary to expectation, these studies find no evidence that flavor-nutrient learning promotes obesity. While of course rodent models do tend to gain weight rapidly and eventually develop obesity when maintained on a highly palatable, energy dense diet, and there are also clear heritable differences in that outcome, there remains little evidence that FNL is a contributing factor. On the contrary, Experiment 3 provided novel evidence that sensitivity to FNL may confer obesity resistance rather than susceptibility.

Experiments 1 and 2 tested whether FNL responses of selectively-bred Obesity-Prone (OP) rats differed from Obesity-Resistant (OR) rats prior to any obesogenic diet exposure, thereby differing only in congenital obesity *proneness* but not in weight or diet history. No strain differences in any measure of FNL were observed in either experiment. Experiment 1 was initially designed to extend our previous work linking FNL to weight gain (Wald & Myers, 2015), in which we maintained rats on an obesogenic high-fat/sugar diet for an extended period and identified the top and bottom third of weight gainers. Comparing each of those groups to chow-fed controls in a FNL paradigm revealed that rats who gained the most weight also showed significantly stronger FNL responses, including in immediate appetition, conditioned preference, and conditioned acceptance, suggesting a positive link between FNL and obesity. However, it could be that inherent pre-existing differences linked to FNL conditionability caused that group to become obese, or that their history of excessive fat/sugar consumption and/or the physiological consequences of weight gain altered their FNL responses. If the former was true, rats known to be obesity prone would be expected to show heightened FNL conditionability while still lean. However, that was not the case. Experiment 2 came to the same conclusion using a different FNL training protocol.

The unexpected and truly novel finding is the initial evidence that FNL may be *protective* against obesity. Experiment 3 screened young adult rats for individual differences in FNL before extended access to a palatable cafeteria diet. Conditioned CS+ acceptance was a significant negative predictor of weight gain. In other words, the tendency to increase intake of a nutrient-paired flavor was associated with obesity resistance. Conditioned CS+ acceptance remained a robust predictor even when controlling for sweet solution intake, and surprisingly, also predicted weight gain independently of a measure of cafeteria diet kcal intake. It should be noted that for practical reasons cafeteria intakes were only measured for the first and last four days of the 24-day diet exposure, and measurements can be imprecise because of rats’ tendency to scatter foods, meaning kcal measurement was only an approximation. Nevertheless, individual differences in conditioned acceptance measured prior to initiating the cafeteria diet served as an independent predictor of weight gain. It is also noteworthy that the FNL measure was not directly correlated with measured cafeteria diet intake, suggesting that if FNL is causally involved in limiting weight gain, it may be acting on pathways other than directly influencing total consumption.

These observations about conditioned acceptance suggest a need to reconsider the functions of FNL in food intake and energy balance, particularly whether FNL acts as a counter-regulatory or a stabilizing influence on body weight. Clearly, as shown by extensive prior work, the behavioral changes produced by FNL all act to increase energy intake in the short term, and there is evidence that FNL does so through both hedonic (liking) and incentive (wanting) aspects of flavor evaluation (reviewed in Berthoud et al., 2021; Myers, 2018). In other words, the conventional view has long been that FNL makes a food “better,” causing it to be preferentially selected and consumed to excess. Operating in an environment with ample energy-dense foods with distinctive flavors and varied macronutrient compositions, FNL would therefore be expected to promote chronic positive energy balance. Some of the earliest electronic esophagus studies demonstrated that when given long-term access to a flavored solution paired with IG nutrient infusion, conditioned acceptance stimulated increased consumption (and therefore nutrient infusion) without fully compensatory reductions in chow intake, increasing total energy intake (Sclafani & Nissenbaum, 1988). However, the current evidence that learning to increase consumption of a nutrient-paired flavor actually predicts resistance to weight gain is at odds with the conventional view. Is increasing energy intake by enhancing liking and/or wanting perhaps not the sole function of FNL?

Another possibility is suggested by Woods’ ‘learned tolerance’ model (Woods, 1991), which emphasizes that associative learning about food-predictive cues enables animals to maximally exploit limited eating opportunities by recruiting adaptive physiological preparatory responses in digestion and metabolism that enable the organism to cope efficiently with large meals consumed rapidly. Increasing intake in FNL does not necessarily reflect the food being perceived as “better,” but rather that conditioning enables a capacity to better handle brief, rapid positive shifts in energy balance.

Considering FNL as a type of Pavlovian conditioning, it seems especially fitting that Pavlov’s foundational work on conditional reflexes focused on learned control of digestive preparation reflexes, not evaluative or motivational responses (Pavlov, 1906). From that perspective, some learned appetition effects may function as cephalic-phase preparatory responses, like the better-known cephalic insulin response that is reflexively elicited by sugar sensing in the mouth but comes under learned control of meal-related conditioned stimuli (Strubbe, 1992; Woods et al., 1977). Some initial support for this view may be seen in another early electronic esophagus experiment in which rats in a FNL paradigm increased their total daily kcal by 13% but did not gain weight (Sclafani et al., 1996), and by another study finding that rats gained more weight on a diet with inconsistent flavor-nutrient relationships (Warwick & Schiffman, 1991). In this view, conditioned acceptance behavior follows from the capacity to make appropriate allostatic adjustments to dietary conditions, and animals who are more capable of these learned adjustments will be more resistant to the dysregulatory effects of an obesogenic food environment. This is by no means contradictory with the view of FNL as increasing liking and/or wanting of the food, as these functions may be complementary. In Experiment 3, conditioned acceptance predicted weight gain but conditioned preference did not, consistent with viewing acceptance as a tolerance-like effect and preference as a hedonic/incentive effect.

Additional work extending the findings of Experiment 3 is needed. First, since most work on the neural and behavioral mechanisms of FNL in rodent models uses the IG infusion paradigm, it will be useful to determine whether the findings of Experiment 3 extend to that paradigm. Conceivably, FNL produced by consuming flavored glucose in the oral training protocol may be uniquely predictive of individual differences in obesity, as it involves learning about the CS flavor but also about the taste of glucose itself, which would be broadly relevant in adapting to a cafeteria diet. In the past decade, clear evidence has emerged that glucose has a discriminable orosensory quality separate from the salient sweet taste mediated by the canonical T1R2/T1R3 ‘sweet’ receptor (Mai et al., 2026; Schier & Spector, 2016). A non-sweet ‘glucose taste’ quality appears to be initially hedonically neutral for naive animals, but may acquire motivational value with experience consuming glucose (Ascencio Gutierrez et al., 2023; Myers et al., 2020). The physiological and behavioral functions of this glucose taste quality are proposed to involve ‘metabolic priming’ functions that contribute to the control of food choice and meal size (Simental-Ramos et al., 2026). Even if the predictive relationship found in Experiment 3 is limited to FNL produced by oral training, that would not make it less important. The IG paradigm offers the advantage of experimental control through complete separation of oral and gut stimuli, but the oral paradigm is arguably a more ecologically valid model of how FNL actually happens in ordinary eating. Therefore, additional work using both paradigms in parallel will be worthwhile. The potential differences between the oral-training and IG-training paradigms could also be the key to explaining the previous contradictory outcomes of FNL after diet-induced obesity (Wald & Myers, 2015; Woods et al., 2016). FNL produced by oral training may be impacted differently by obesity because of how diet history impacts perceiving or responding to the non-sweet, metabolic priming functions of oral glucose sensing.

A related question concerns whether the association between conditioned acceptance and subsequent weight gain is specifically tied to FNL with glucose or would extend to conditioning with other macronutrients. Glucose is maximally effective at generating appetition signals and producing FNL, especially compared to fructose, which is almost entirely ineffective (Berthoud et al., 2021; Zukerman et al., 2013). FNL also occurs with fats (McDougle et al., 2024; Zukerman et al., 2013) and with proteins or amino acids (Ackroff & Sclafani, 2016; Murphy et al., 2018), but learning may be weaker or slower than for glucose (Ackroff et al., 2009; Myers, 2013; Revelle & Warwick, 2009), and individual differences have not been explored. Further, as previously described, learned responses to glucose may also recruit a glucose-specific oral sensing pathway. Given this evidence that mechanisms of FNL are at least partly macronutrient-specific, and also that intakes of macronutrients are differentially defended (Berthoud et al., 2012; Raubenheimer & Simpson, 2023), additional work should explore whether the conditioned acceptance effect with fat-paired or protein-paired flavors also predicts resistance to cafeteria weight gain, and further, whether the predictive relationship between conditioned acceptance and weight gain depends on the macronutrient composition of the obesogenic diet. It is unclear from the current data whether conditioned acceptance for a glucose-paired flavor predicted rats’ responses to the cafeteria diet generally, or specifically to its glucose content.

The current conclusions are also limited by including only female rats. The degree to which FNL itself is sex-dependent in rodent models is not widely studied, as both males and females have been used in FNL experiments but few studies have directly compared sexes within-experiment (e.g., Nyema et al., 2023). Sex differences are documented in other types of food-based associative learning, for example, with females exhibiting stronger palatable food-induced conditioned place preference (Sinclair et al., 2017) and faster acquisition but slower extinction of Pavlovian appetitive conditioning (Lafferty et al., 2026; Pitchers et al., 2015). Sex differences in patterns of diet-induced obesity are also evident, with female rodents showing less hyperphagia and slower weight gain than males when fed high-fat diets (Huang et al., 2020; Maric et al., 2022). Whether the inverse relationship between FNL and weight gain observed here is also seen in males, and whether differences in learning play a role in the sex differences in weight gain, will need additional study.

In conclusion, although these experiments address only some potential inter-relationships between FNL and obesity, they do challenge the premise that the well-known ability of FNL to increase short-term energy intake in rodent models makes FNL a causal factor in long-term overconsumption leading to obesity, and instead suggest FNL may protect against obesity. A promising avenue for translational research therefore could be concerned with how aspects of the modern obesogenic environment impact this apparent protective function. The metabolic ‘tolerance’ perspective on FNL could potentially inform the long-standing conundrum over why evidence for *de novo* FNL in well-controlled human experiments has been elusive, at least as measured by self-reported liking and/or wanting or other behavioral measures of “value” (see reviews by Brunstrom, 2005, 2007; Yeomans, 2012). Studies incorporating functional brain imaging have since observed that FNL establishes distinct conditioned responses in the nucleus accumbens and hypothalamus that are largely unrelated to shifts in flavor liking, and instead reflect shifts in glucose homeostasis experienced during the FNL training trials (de Araujo et al., 2013). Another recent human study has documented an association between individual differences in FNL and markers of metabolic health. Participants who showed better behavioral discrimination between a nutrient-paired CS+ and CS− flavor also tended to have lower fasting glucose and HbA1c (Baugh et al., 2025). Collectively, this work suggests that understanding relationships between FNL, diet, and weight in both rodent models and humans could be informed by incorporating a focus on metabolic responses in addition to changes in flavor evaluation.

## Research Data

Copies of data files with analysis code and output (JASP) will be available at https://osf.io/p5h6z upon publication.

## Statement on Generative AI use

This paper was written by the author using his own, organic brain. No part of the manuscript was generated by AI (with the possible exception of changes introduced by the publisher’s AI copyediting and page layout tools, which are beyond the author’s control). This paper is intended to be read, interpreted, and discussed by corporeal, sentient beings using their own intellectual faculties.

## Acknowledgements

This paper is an invited contribution in conjunction with receiving the Hoebel Award for Creativity at the 2025 SSIB meeting. I would like to express my sincere gratitude, especially for the outstanding training and guidance I received from my mentors Tim Cannon (University of Scranton), Ted Hall (Duke) and Tony Sclafani (Brooklyn College-CUNY), the support of my scientific big sisters Susie Swithers and Karen Ackroff, and for the encouragement of many friends, collaborators, and informal mentors in SSIB who are too numerous to list. The experiments reported here were conducted with the invaluable assistance of several talented Bucknell undergraduate students including Marta Majewski, Hannah Goldberg, Zach Mahaney, Meredith Lutz, Julia Mammone, Kristen Palframann, and Peter Lanzi. I am grateful to Dr. Carrie Ferarrio for supplying the OP/OR rats, and for the outstanding assistance of Cindy Rhone and the Bucknell animal caretaking staff.

