## Supplemental Methods for "Does flavor-nutrient learning promote or protect against diet-induced obesity? Individual differences in conditionability predict resistance to weight gain in rats"

Supplementary Methods

*Cafeteria diet*

|  | |  | | kcal/g | | fat kcal/g | | carb kcal/g |
| --- | --- | --- | --- | --- | --- | --- | --- | --- |
| 1 | Sugar/fat whip 70/20/0/10 (rum flavor) | | 4.6 | | 1.8 | | 2.8 | |
|  | Wheat breakfast cereal (Total, General Mills) | | 3.5 | | 0.2 | | 2.5 | |
|  | Potato chips (Utz brand original unsalted) | | 5.4 | | 2.9 | | 1.9 | |
| 2 | Sugar/fat whip 56/15/11/18 (almond) | | 3.6 | | 1.4 | | 2.2 | |
|  | Parmesan crackers (Pepperidge Farm) | | 4.7 | | 1.5 | | 2.5 | |
|  | Vanilla creme wafer cookies (Great Value) | | 5.0 | | 1.9 | | 2.8 | |
| 3 | Sugar/fat whip 42/12/16/20 (hazelnut) | | 2.8 | | 1.1 | | 1.7 | |
|  | Honey graham breakfast cereal (Malt-O-Meal) | | 4.3 | | 0.8 | | 2.8 | |
|  | Fried pork rinds (Utz brand) | | 5.7 | | 3.6 | | 0.3 | |
| 4 | Custard dessert (Lucky Leaf, root beer flavor) | | 1.5 | | 0.6 | | 0.8 | |
|  | Chocolate breakfast cereal (Malt-O-Meal) | | 4.0 | | 0.5 | | 3.3 | |
|  | Hot dog (Bar S brand beef franks) | | 2.9 | | 2.1 | | 0.4 | |
| All days: | Chow (LabDiet 5001) | | 3.4 | | 0.6 | | 2.6 | |

Cafeteria diet consisted of twelve different foods grouped into four trios. A different trio was given each day along with ad libitum chow, cycling through all four trios in each four-day period in semi-random order. Therefore, each trio of cafeteria foods was given six times in 24 days. The foods given each day were provided ad libitum in separate containers hung inside the home cage. For the sugar/fat whips, numbers indicate percentage by weight of sucrose, vegetable shortening cellulose, and water, with an added flavor extract added (McCormack brand, 0.5%).
